# Predation-resistant *Pseudomonas* bacteria engage in symbiont-like behavior with the social amoeba *Dictyostelium discoideum*

**DOI:** 10.1101/2023.10.02.560472

**Authors:** Margaret I. Steele, Jessica Peiser, Joan E. Strassmann, David C. Queller

**Affiliations:** Biology Department, Washington University in St. Louis, St. Louis, Missouri, USA

## Abstract

The soil amoeba *Dictyostelium discoideum* acts as both a predator and potential host for diverse bacteria. We tested fifteen *Pseudomonas* strains that were isolated from transiently infected wild *D. discoideum* for ability to escape predation and infect *D. discoideum* fruiting bodies. Three predation-resistant strains frequently caused extracellular infections of fruiting bodies but were not found within spores. Furthermore, infection by one of these species induces secondary infections and suppresses predation of otherwise edible bacteria. Another strain can persist inside of amoebae after being phagocytosed but is rarely ingested. We sequenced isolate genomes and discovered that predation-resistant isolates are not monophyletic. Many *Pseudomonas* isolates encode secretion systems and toxins known to improve resistance to phagocytosis in other species, as well as diverse secondary metabolite biosynthetic gene clusters that may contribute to predation resistance. However, the distribution of these genes alone cannot explain why some strains are edible and others are not. Each lineage may employ a unique mechanism for resistance.

## Introduction

Most eukaryotic organisms interact with bacterial symbionts throughout their lives. Some symbionts are vertically transmitted, such as the *Buchnera* endosymbionts of aphids (Douglas, 1998), while others are acquired from the environment, such as the bioluminescent *Vibrio fischeri* symbionts of bobtail squid (Visick et al., 2021) or nitrogen fixing rhizobia of legumes (Mithöfer, 2002). These established symbioses are often facilitated by complex adaptations, such as specialized host structures for housing mutualistic symbionts. Similarly, parasitic symbionts often alter and exploit host structures to facilitate their own reproduction, as is seen in pathogens such as *Legionella pneumophila* and *Mycobacterium tuberculosis* that can replicate within phagocytic cells (Queval et al., 2017; Swanson & Hammer, 2000). However, these complex interactions likely originated from much simpler chance encounters or opportunistic infections. Predatory amoebae could be a source of novel intracellular pathogens because they constantly interact with bacteria in the environment and select for traits that contribute to survival of phagocytosis (Molmeret et al., 2005). Following the same logic, amoebae may provide an opportunity to study the early stages of the evolution of mutualistic and parasitic bacterial symbionts.

The soil amoeba *Dictyostelium discoideum* is a simple, tractable organism that interacts with bacteria in different ways throughout its lifecycle. *D. discoideum* spends much of its life as a unicellular amoeba, consuming bacteria through phagocytosis and replicating via binary fission. Although *D. discoideum* is capable of preying upon a diverse range of bacteria, predation resistant bacteria have been identified in multiple phyla (Brock et al., 2018). Little is known about the mechanisms these species use to escape predation. When *D. discoideum* amoebae run out of prey, tens to hundreds of thousands of amoebae aggregate to form a motile, multicellular slug. A fraction of these cells become sentinel cells, which eliminate potential bacterial pathogens by sequestering them within vacuoles and producing extracellular traps (Brock et al., 2016; Chen et al., 2007). The slug migrates towards the soil surface, where it develops into a fruiting body that consists of a spore-filled sorus supported by a stalk, which aids in dispersal of spores by insects (smith et al., 2014). The sorus is typically free from bacteria. Dispersed spores hatch into amoebae when prey bacteria are present (Shu et al., 2021). Bacteria capable of surviving phagocytosis, escaping the neutrophil-like activities of sentinel cells, and infecting the sorus may benefit from co-dispersing with spores or preying upon spores or hatched amoebae.

In addition to preying upon bacteria, *D. discoideum* acts as a host for both beneficial and pathogenic bacteria. *Paraburkholderia agricolaris*, *Paraburkholderia hayleyella*, and *Paraburkholderia bonniea* are conditionally beneficial intracellular symbionts of *D. discoideum* that may be acquired from the environment or vertically inherited. These bacteria are inedible themselves but induce secondary infections by edible bacteria, allowing *D. discoideum* spores to co-disperse with prey bacteria that can seed new populations (DiSalvo et al., 2015; Scott et al., 2022). Infection by *Paraburkholderia* can therefore be beneficial to spores that disperse to areas with limited prey. *D. discoideum* amoebae can also be infected by numerous human intracellular pathogens under laboratory conditions, making it a popular model system for studying bacterial pathogenesis (Cardenal-Muñoz et al., 2018; Clarke, 2010; Cosson & Lima, 2014; Dunn et al., 2018). In nature, amoebae sometimes act as environmental reservoirs for pathogenic bacteria such as *Bordetella bronchiseptica* (Taylor-Mulneix et al., 2017) and *Mycobacterium bovis* (Butler et al., 2020). However, far more is understood about the mechanisms human pathogens use to infect *D. discoideum* than is known about the bacteria it interacts with in nature.

A recent study of bacteria isolated from wild *D. discoideum* showed that many soil bacteria form short-lived associations with *D. discoideum* (Brock et al., 2018). Many of these isolates belonged to genus *Pseudomonas*, including both edible and predation resistant strains. To better understand the evolution of predation resistance and pathogen-like behaviors in soil *Pseudomonas* species, we tested fifteen *Pseudomonas* isolates for susceptibility to predation and ability to infect the sorus. These strains were isolated from wild *D. discoideum* and are therefore known to interact with the predator in nature. We used a combination of genome sequencing, microscopy, and infection assays to explore the evolution of predation resistance in soil *Pseudomonas* species and to look for parallels between the effects of predation-resistant *Pseudomonas* and symbiotic *Paraburkholderia*.

## Methods

### Strains, media, and culture conditions

Bacterial strains, *D. discoideum* clones, and plasmids are listed in Table S1. Antibiotics and additives were used at the following concentrations, except where indicated: 20 µg/ml gentamicin (Gm), 30 µg/ml kanamycin (Km), 100 µg/ml carbenicillin (Crb), 300 µg/ml streptomycin (Str), 60 µg/ml spectinomycin (Spec), 20 µg/ml chloramphenicol (Cm), 0.3 mM 4,6-diaminopimelic acid (DAP). *Klebsiella pneumoniae* (Kp) and *Pseudomonas* spp. were grown on LB (Fisher) plates at 30°C. *Escherichia coli* strains used for conjugation were grown in LB broth with appropriate antibiotics at 37°C, shaking at 225 rpm. Axenic *D. discoideum* cultures were incubated at 21°C, shaking at 100 rpm in 20ml HL5 medium including glucose (Formedium) supplemented with Crb and Str to prevent bacterial contamination. For *D. discoideum* experiments on agar, 4×10^5^ spores and 200µl of a bacterial suspension were spread on SM/5 agar plates ( 2 g glucose (Fisher Scientific), 2 g BactoPeptone (Oxoid), 2 g yeast extract (Oxoid), 0.2 g MgCl_2_ (Fisher Scientific), 1.9 g KHPO_4_ (Sigma-Aldrich), 1 g K_2_HPO_5_(Fisher Scientific), and 15 g agar (Fisher Scientific) per L). Bacterial suspensions were prepared by diluting bacteria collected from LB plates to an OD_600_ of 1.5 in KK2 ( 2.25 g KH _2_PO_4_ (Sigma-Aldrich) and 0.67 g K_2_HPO_4_ (Fisher Scientific) per L). *D. discoideum* cultures on agar were incubated at room temperature (20-22°C).

### Quantification of bacterial CFU

Bacteria were collected from liquid or by flooding agar with 1 ml of KK2. Seven 1:10 serial dilutions were prepared and 10 µl droplets of each dilution were spotted in triplicate on agar. Plates were incubated at 30°C and CFU were counted after 1-2 d.

### Chromosomal insertion of fluorescent protein genes

Site-specific integration of *gfp* and antibiotic resistance markers into genomes of *Pseudomonas* isolates was achieved using the mini-Tn7 system, as previously described (Choi & Schweizer, 2006). Briefly, *E. coli* M F D *pir*, a DAP auxotroph, was used to deliver plasmids carrying a transposon and transposase genes (see Table S1) to recipient cells through conjugation. Equal numbers of donor, helper, and recipient cells were mixed and spotted on LB DAP plates, then incubated at 30°C overnight. Cells were collected, washed with KK2, and then spread on selective LB plates. Kp was labeled with E2-crimson using the same method.

### Edibility assays

To classify *Pseudomonas* strains as edible or inedible, *D. discoideum* QS157 spores were spread on SM/5 agar plates with *Pseudomonas* sp. as a sole food source or with a 10:90 mixture of *Pseudomonas* sp. and Kp. As controls, QS157 was also grown on Kp, *Pa. bonniea* Bb859 (a slightly edible symbiont species), and *Ps. aeruginosa* PAO1 (an inedible pathogen). Plates were monitored for 7 d for fruiting body development. After 7 d, cells were collected to quantify the remaining bacterial CFU.

### Bacterial carriage assays and CFU per sorus

To determine whether *Pseudomonas* strains infect the sorus of *D. discoideum* fruiting bodies, *D. discoideum* QS157 spores were spread on SM/5 plates with either 100% *Pseudomonas*, 50% *Pseudomonas* and 50% Kp, or 10% *Pseudomonas* and 90% Kp. Individual sori were collected after 7 d and spotted onto the surface of SM/5 agar plates and the number of spots that showed bacterial growth was recorded to determine the fraction infected by bacteria. To enumerate the bacteria per sorus, individual sori were transferred to PCR tubes containing 200 µl of KK2 supplemented with 0.05% NP-40 alternative. Tubes were vortexed to release spores and bacteria from sori, then bacterial CFU were quantified.

### Intracellular survival assay

To determine how long bacteria survived after phagocytosis, we used an intracellular survival assay similar to that described by Pukatzki et al. (Pukatzki et al., 2002). Axenically grown *D. discoideum* AX4 amoebae were washed three times with cold KK2. Washed amoebae were resuspended in HL5 at a concentration of 2×10 ^6^ cells/ml. Pv, Pl, Pp, *Ps. aeruginosa* PAO1, Kp, and *Pa. bonniea* Bb859 were grown overnight in LB broth at 30°C, 225 rpm, then spun down and resuspended in HL5 at OD_600_ 1.

1 ml volumes of amoebae were transferred to wells of two 24-well tissue culture plates. After 1.5 h, 50µl of bacteria were added to wells. The 24-well plates were centrifuged for 10 min at 750 rcf, then incubated at room temperature for 30 min. The supernatant was removed, leaving only attached cells, 1 ml of KK2 was added and then removed to wash the cells, then 1 ml of KK2 with 400 µg/ml gentamicin was added to each well to kill extracellular bacteria. Plates were left at room temperature for either 3 or 22 h to determine how long bacteria were able to survive within amoebae. The number of intracellular bacteria was determined by collecting the cells from each well and washing them three times with cold KK2 to remove the gentamicin.

Cells were finally resuspended in 1 ml of KK2 with 0.05% Triton-X 100 to lyse the amoebae. Bacterial CFU were then quantified to determine the number of intracellular bacteria recovered from each well. The time between addition of gentamicin and lysis of amoebae was approximately 5 and 24 h for the two timepoints.

### Protection of other species

Three edible strains, Ph-GmR-GFP (14P 8.1_Bac3), Pe-KmR-GFP (7P 10.2_Bac1), and Kp-GmR-GFP, and three inedible strains, Pv, Pl, and Pp, were grown overnight at 30°C, 225 rpm in SM/5 broth, then diluted to OD _600_ 1. Edible strains were mixed 1:1 with inedible strains. Controls without predation-resistant bacteria were mixed with sterile broth. *D. discoideum* AX4 amoebae were grown axenically, then washed and resuspended in SM/5 broth at a concentration of 2×10^6^ cells/ml. 65 µl of bacteria and 15 µl of amoebae were spread on 60 mm SM/5 agar plates and incubated at room temperature. After 7 d, all cells on the plate were collected and bacterial CFU were quantified to determine the number of edible and inedible bacteria remaining.

### Microscopy

The contents of sori infected with GFP-labeled *Pseudomonas* strains were examined using confocal microscopy to determine whether bacteria in the sorus are intracellular. *D. discoideum* was grown with *Pseudomonas* (10%) and Kp, as described above. Sori were collected from fruiting bodies and suspended in 50 µl KK2 with 1% calcofluor. 10 µl was placed on top of a 1% agarose pad on a microscope slide, prepared by using a siliconized glass cover slip to flatten a 125 ml drop of agarose, then covered with a cover slip.

To visualize interactions between *Pseudomonas* and amoebae, microscope slides were embedded in petri plates under a thin layer of 0.5% agar SM/5. GFP-labeled *Pseudomonas* strains and Kp were suspended in KK2 at OD 1.5. Each *Pseudomonas* strain was mixed with Kp at a 1:1 ratio, then 200µl of the mixture and 4×10 ^5^ QS9-mCherry spores were spread on agar over embedded slides. After approximately 42 h, slides were cut out of the agar plate, a cover slip was placed on top of the agar, and cells were imaged.

### Genome and 16S rRNA gene sequencing

Genomic DNA was extracted from bacteria using a Qiagen DNeasy blood and tissue kit. 10µl of DNase-free RNase (MilliporeSigma) was added to each sample after Proteinase K treatment to remove RNA. 16S ribosomal RNA genes of *Pseudomonas* soil isolates were sequenced to verify that all isolates belonged to genus *Pseudomonas* and to identify closely related reference genomes. 16S ribosomal RNA genes were amplified through PCR using universal 16S primers 27F (5’-AGAGTTTGATCCTGGCTCAG-3’) and 1507R (5’-TACCTTGTTACGACTTCACCCCAG-3’). We used a touchdown PCR program with an initial annealing temperature of 60°C, which was reduced by 1°C each cycle for the first 10 cycles, followed by 30 cycles with an annealing temperature of 50°C. PCR products were purified and sequenced (Azenta Life Sciences). Related reference genomes were identified using nucleotide BLAST (Camacho et al., 2009) to search the NCBI 16S ribosomal RNA sequences database. Genomes were sequenced by the Microbial Genome Sequencing Center (MiGS) using an Illumina NextSeq 2000 to generate 2×151 bp paired end reads. 1.3 to 1.7 million reads were obtained for each genome, providing 56 to 96x coverage.

### Genome assembly and annotation

FastQC 0.11.9 (Andrews, 2010) was used to check quality of reads. Reads were assembled using Unicycler v0.4.9 (Wick et al., 2017) and Quast 5.1.0rc1 (Gurevich et al., 2013) was used to assess the quality of the assemblies. The assembled genomes were annotated using the Rapid Annotation through Subsystem Technology (RAST) platform (Aziz et al., 2008) and then re-annotated using the NCBI Prokaryotic Genome Annotation Pipeline (Tatusova et al., 2016). AntiSMASH (Blin et al., 2021; Medema et al., 2011) was used to identify clusters of genes encoding secondary metabolite biosynthesis pathways. These clusters were grouped using all-v-all BLAST to identify sequences that shared ≥70% nucleotide identity over ≥20% of the query length. Groups were visualized using Cytoscape v3.9.0 (Su et al., 2014).

To determine whether the *Pseudomonas* genomes encode Type III secretion systems (T3SS), representative protein sequences for T3SS structural proteins SctJNQRSTUV, which are found in both T3SS and flagella; SctC, found only in T3SS; and FlgBC and FliE, found only in flagella (Abby & Rocha, 2012), were downloaded from the NCBI protein database. BLAST+ v2.9.0 was used to search for homologous sequences within a local BLAST database containing proteins encoded by the *Pseudomonas* genomes. To identify Type VI secretion systems (T6SS), reference sequences for 13 structural proteins (TssA-TssM) were downloaded from the SecReT6 database (Li et al., 2015). CD-HIT v4.8.1 (Fu et al., 2012) was used to cluster sequences that shared more than 40% amino acid identity, and then reference sequences from each cluster were used to search for homologous sequences in the *Pseudomonas* genomes using protein BLAST. T3SS and T6SS in different genomes were grouped based on homology into three distinct T3SS and five T6SS. ExoU, ExoY, ExlA, and MgtC homologs were identified by using BLAST to search all proteins encoded by the *Pseudomonas* genomes for homology to reference sequences from *Ps. aeruginosa* (accessions ASM94169.1, PWU33926.1, QDL04633.1, and BAQ2388.1).

### Genome phylogenies and average nucleotide identity

Orthofinder v2.5.4 (Emms & Kelly, 2017, 2018, 2019) was used to construct species phylogenies based on all single copy orthologs found in all *Pseudomonas* genomes. Reference genomes were selected based on the best matches to the 16S rRNA genes. Phylogenies were visualized using the interactive Tree of Life (iTOL) v6 (Letunic & Bork, 2021) and were further modified using Adobe Illustrator. To determine whether sequenced genomes belong to species with reference genomes in NCBI databases, Average Nucleotide Identity for each species pair was calculated using FastANI (Jain et al., 2018).

### Statistical tests

Fisher’s exact test was performed in R v4.2.0 (R Core Team, 2014). Other statistical tests were performed using GraphPad Prism v9.5.1 (GraphPad Software, San Diego, California USA, www.graphpad.com).

## Results

### Predation-resistant *Pseudomonas* strains are not monophyletic

Some *Pseudomonas* strains are resistant to predation by *D. discoideum*, while others are susceptible. However, it is not known how much diversity exists among predation-resistant strains or whether mechanisms of predation resistance have evolved more than once within the genus. To explore these questions, we selected fifteen *Pseudomonas* strains that were isolated from wild *D. discoideum* clones in two previous studies (Brock et al., 2018; Stallforth et al., 2013). Eight strains were susceptible to predation, meaning *D. discoideum* amoebae were able to clear agar plates of bacteria and form fruiting bodies even when no additional food bacterium was provided. The other seven strains were inedible, with large amounts of bacteria remaining on agar plates and few or no fruiting bodies after 7 d. We tested for the presence of bacteria in the sorus by collecting individual sori from fruiting bodies and transferring them to fresh agar plates. Because *D. discoideum* employs multiple mechanisms to eliminate bacteria during development to protect spores from potential pathogens, the sorus is typically expected to be bacteria-free. However, multiple *Pseudomonas* strains were able to infect the sorus (Fig. 1). We chose to focus on three predation-resistant strains. Strains 20P 3.2_Bac4, 20P 3.2_Bac5, and 18P 8.2_Bac1 infected more often than average (Fisher’s exact test, p < 0.05, FDR correction for multiple comparisons (Table S2). However, 20P 3.2_Bac4 and 20P 3.2_Bac5 are very closely related, so we replaced the latter with the nearly significant 6D 7.1 Bac1, leaving us with 20P 3.2_Bac4, 18P 8.2_Bac1, and 6D 7.1_Bac1, which will be referred to hereafter as Pv, Pp, and Pl. Interestingly, the ability of Pl to infect the sorus is affected by temperature. When *D. discoideum*, Pl, and Kp are co-cultured at 18.6°C, almost all sori are infected by Pl, while no sori are infected at 25°C (Table S3), so it is possible that fluctuations in laboratory temperature reduced the observed frequency of infection.

**Figure 1.**
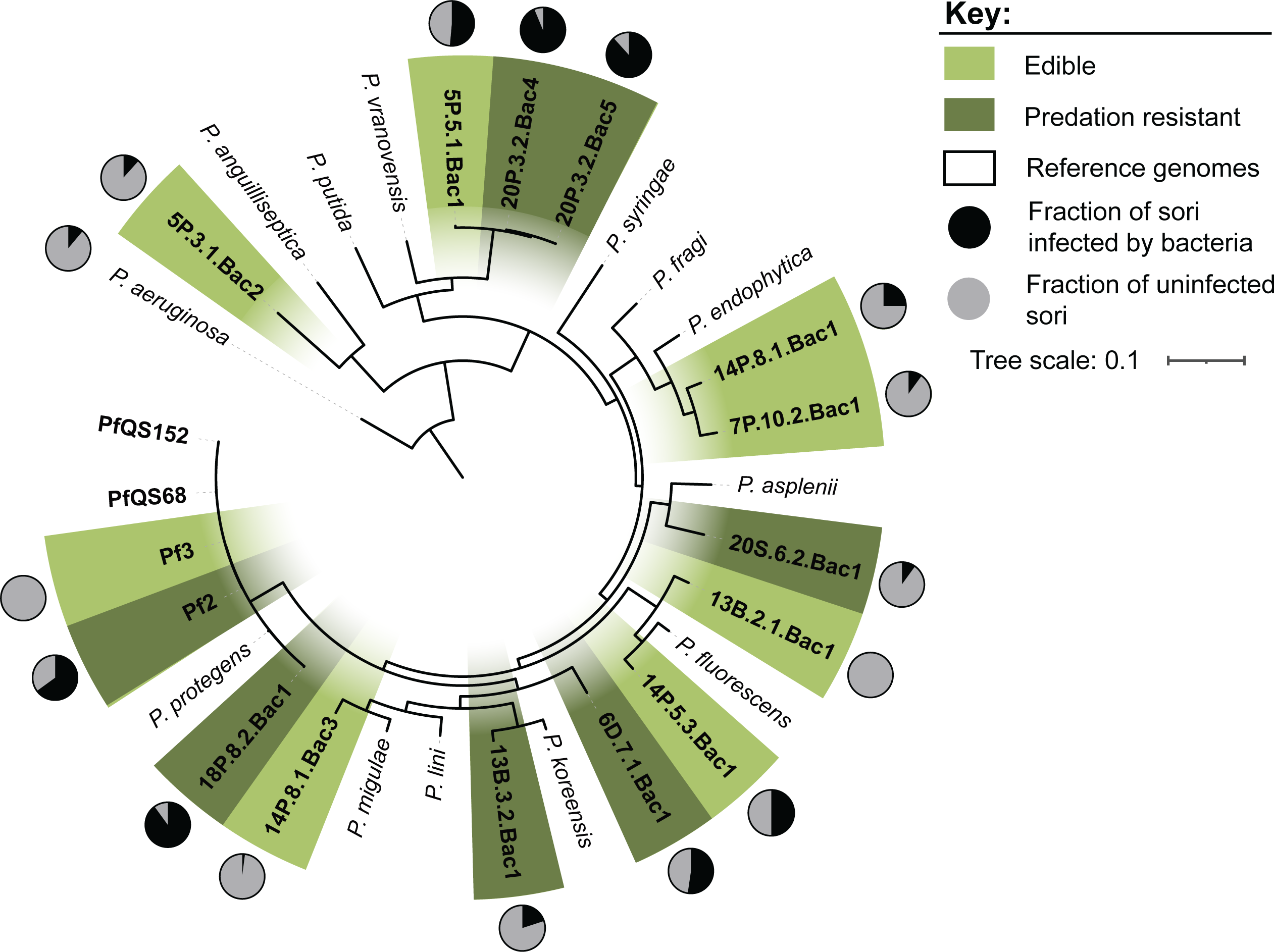
Diverse *Pseudomonas* strains evade predation and infect the sorus of *D. discoideum* fruiting bodies. A phylogeny, based on 2,287 shared single copy proteins, shows the evolutionary relationships between edible *Pseudomonas* isolates (light green), inedible isolates (dark green), and reference genomes (white). Pie charts around the outer edge of the phylogeny show the fraction of sori that were infected with bacteria after *D. discoideum* was grown on a mixture of *Pseudomonas* and *K. pneumoniae*. 10-80 individual sori were sampled for each strain.

We sequenced the genomes of 13 *Pseudomonas* isolates. Two *P. protegens* genomes (Pf2 and Pf3) were previously sequenced. We constructed a phylogeny based on the 15 isolate genomes and reference genomes of related species (Fig. 1). Although all 15 strains belong to the *Pseudomonas fluorescens* complex, the predation resistant strains are not monophyletic. We used Average Nucleotide Identity (ANI) to determine whether each isolate belongs to the same species as the reference genome with the most similar 16S rRNA gene sequence (Table S3). For most isolates, ANI was less than 95% for their most closely related reference genome, suggesting they represent novel species.

### *Pseudomonas* infections of *D. discoideum* are extracellular

To better understand how Pv, Pl, and Pp behave within the sorus, we quantified the number of bacteria per sorus in *D. discoideum* fruiting bodies grown on a mixture of *Pseudomonas* and *Klebsiella pneumoniae* (Kp). Kp is a food bacterium included to facilitate *D. discoideum* growth and fruiting body formation in the presence of predation-resistant *Pseudomonas*. Although not all sori become infected by *Pseudomonas* (Fig. 2A), the number of bacteria within infected sori is consistent, suggesting that bacterial replication within the sorus may be restricted by limited space or nutrient availability (Fig. 2B). The number of *Pseudomonas* cells per sorus is also similar across species, with an average of 2.67×10^5^ Pv, 5.74×10^5^ Pl, and 5.41×10 ^5^ Pp bacteria per sorus. This is significantly less than the average number of *Pa. bonniea* Bb859 cells per sorus, 1.72×10 ^6^. *Pa. bonniea* is an intracellular symbiont that is present inside and outside of spores and has smaller cells and a reduced genome, which may help it replicate to higher densities within sori. As expected, Kp does not infect the sorus on its own. Unlike other *Pseudomonas* species, the presence of Pv frequently allows secondary infections of the sorus by Kp (Fig. 2C,D), a characteristic that is usually associated with symbiotic *Paraburkholderia*. Occasionally, sori of fruiting bodies grown with Pv contain only Kp, suggesting Pv is not required for Kp to survive within the sorus but instead has an effect earlier in development, possibly by inhibiting phagocytosis of Kp.

**Figure 2.**
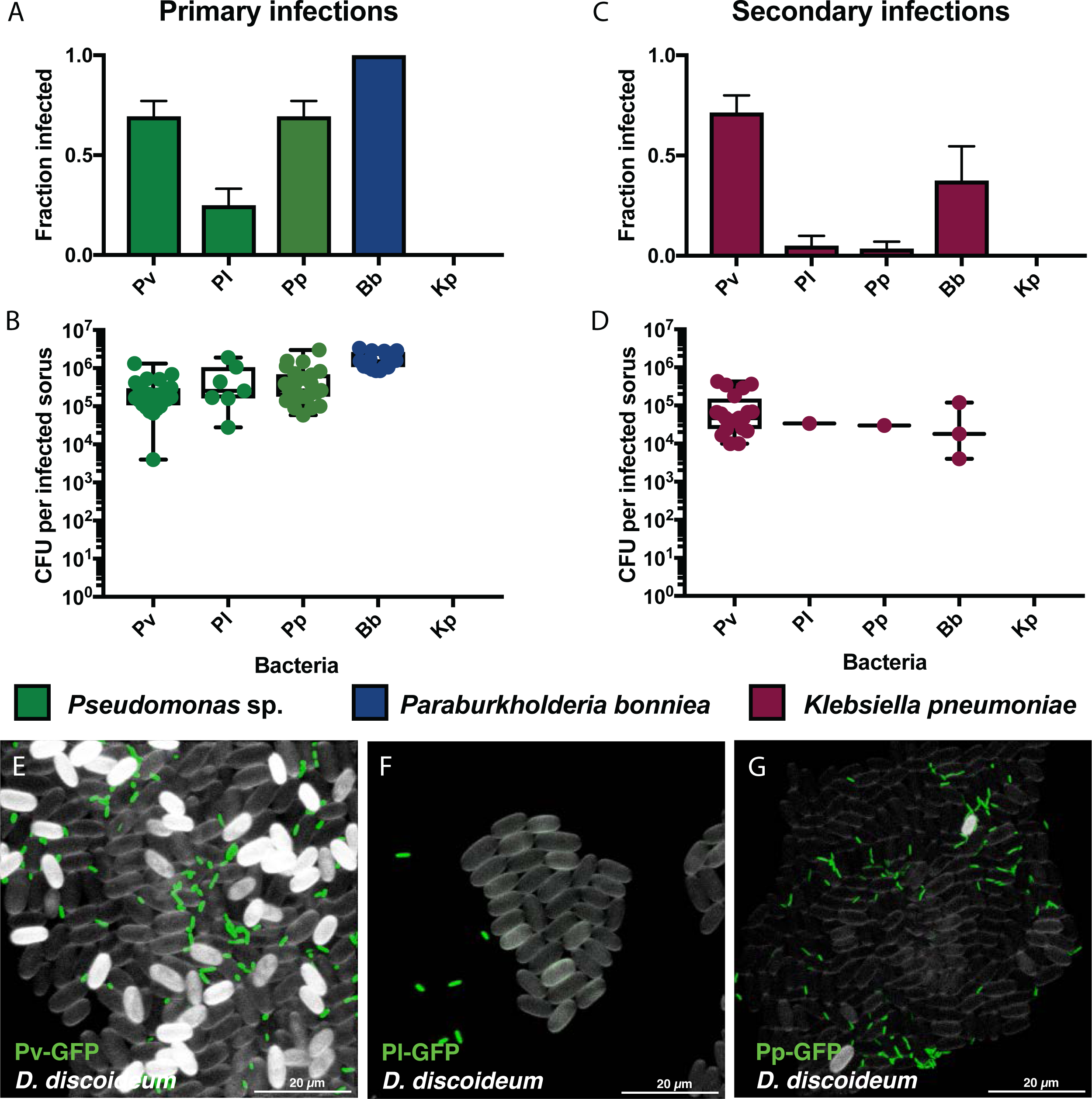
*Pseudomonas* infections within the sorus are extracellular. Sori were collected from *D. discoideum* fruiting bodies grown on mixtures of Kp-E2crimson and GFP-labeled *Pseudomonas* sp., *Pa. bonniea* Bb859 (Bb), or Kp. (A,C) Fraction of sori infected with bacteria. Error bars show standard error. (B,D) Number of bacteria per infected sorus. Each point represents CFU recovered from one sorus. (A,B) Primary infection by *Pseudomonas* isolates or Bb. (C,D) Secondary infection of the same sori by Kp. (E-G) Microscopy images of spores and bacteria collected from sori infected with (E) Pv-GFP, (F) Pl-GFP, and (G) Pp-GFP. Spore coats were stained with calcofluor, shown in white, while bacteria are shown in green.

We used fluorescence microscopy to determine whether bacteria were present inside or outside of spores. None of the three *Pseudomonas* species were found within spores, suggesting that infections within the sorus are extracellular (Fig. 2E-G). The bacteria appear to aggregate with clumps of spores, but cells were removed from the sorus and stained prior to imaging, so images may not accurately depict their spatial organization in situ.

A hallmark of intracellular pathogens is the ability to infect and replicate within host cells. To determine whether Pv, Pl, or Pp replicate within amoebae, we cocultured each strain with axenically-grown AX4 amoebae, added gentamicin to eliminate extracellular bacteria, then incubated the cells for 3 h or 22 h to determine how long each strain persisted intracellularly.

We included *Pseudomonas aeruginosa* PAO1, Kp, and *Pa. bonniea* Bb859 for comparison, with the expectation that PAO1, a pathogen, and *Pa. bonniea*, an intracellular symbiont, would persist after phagocytosis, while Kp would be rapidly ingested and eliminated. After 3 h of gentamicin treatment, we detected an average of 1.00×10^2^ Pv, 2.09×10 ^4^ Pl, 3.14×10 ^5^ Pp, 2.02×10^4^ PAO1, 2.39×10 ^6^ Kp, and 8.22×10 ^6^ *Pa. bonniea* bacteria per 2×10 ^6^ amoebae (Fig. 3). These counts suggest that most amoebae in the Kp and *Pa. bonniea* treatment groups had ingested bacteria, at least some of which were still alive after 3 h. In contrast, only 5 in 100 000 amoebae contained Pv, 1 in 100 contained Pl, and 1.6 in 10 contained Pp. After 22 h, Pp (3.52×10^3^ bacteria per 2×10 ^6^ amoebae) and *Pa. bonniea* (2.84×10 ^4^ bacteria) were still present inside amoebae, but the other species were below the limit of detection. We also detected a small amount of PAO1 in one replicate. *Pa. bonniea* Bb859 is known to be slightly more edible than other symbiotic *Paraburkholderia* (Haselkorn et al., 2019), which may explain the decrease in the number of intracellular bacteria over time.

**Figure 3.**
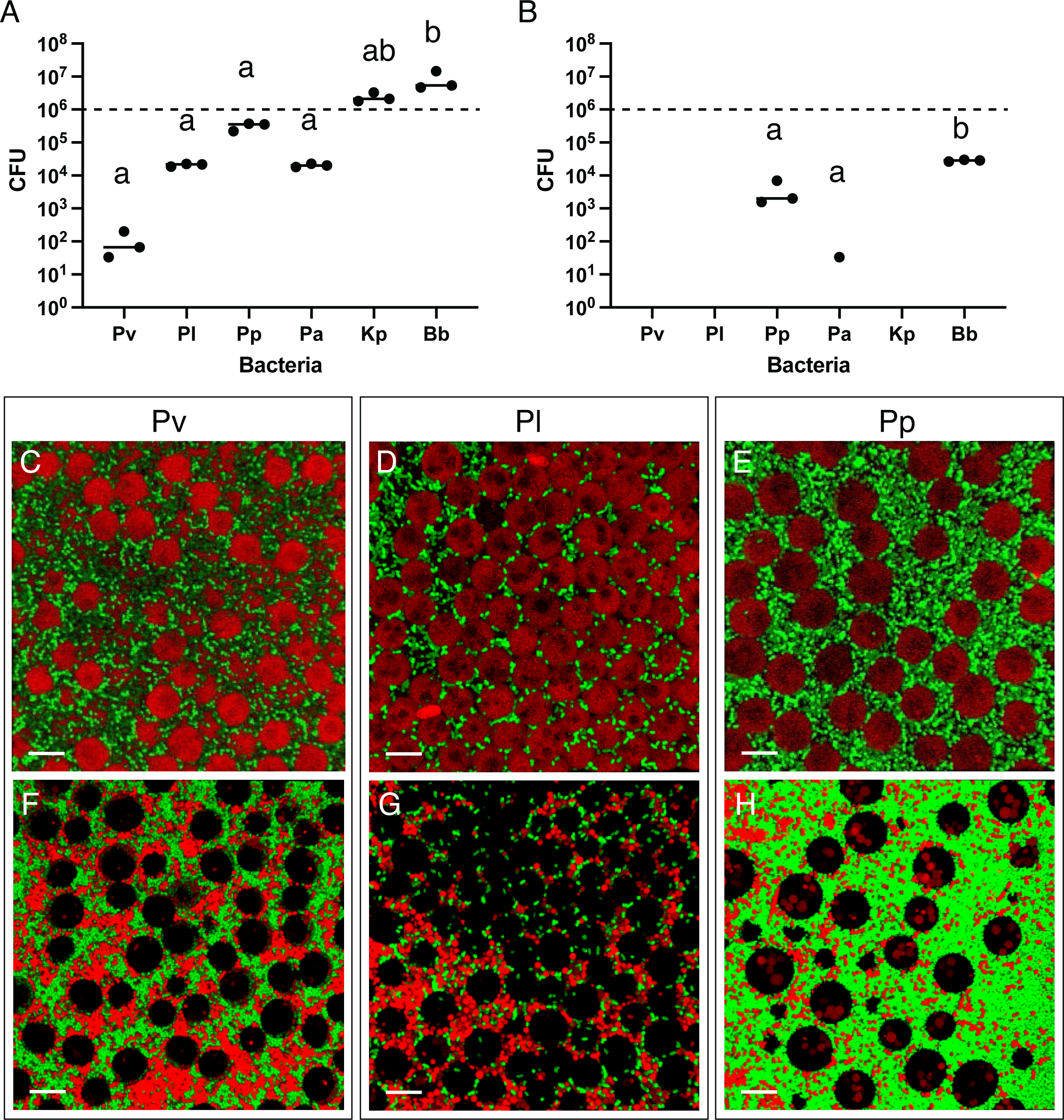
*Pseudomonas* sp. infections of amoebae are uncommon. Mixtures of *D. discoideum* AX4 amoebae and bacteria ( *Pseudomonas* isolates Pv, Pl, and Pp; Pa, pathogen *Ps. aeruginosa* PAO1; Kp, food bacterium *K. pneumoniae*; and Bb, intracellular symbiont *Pa. bonniea* Bb859) were treated with gentamicin to eliminate extracellular bacteria. After 3 h (A) or 22 h (B), amoebae were washed and lysed. Points show bacterial CFU recovered from lysate, while the dashed line represents the number of amoebae. Different letters indicate p ≤ 0.05, one-way ANOVA with Tukey’s multiple comparisons correction. Confocal microscopy was used to image bacteria and amoebae on soft agar to determine whether bacteria were located within amoebae. (C) Pv-GFP, (D) Pl-GFP, and (E) Pp-GFP were grown with *D. discoideum* QS9.1-mCherry and unlabeled Kp (C-E) or with *D. discoideum* QS9.1-mCherry and Kp-E2crimson (F-H) on agar.

To verify that inedible *Pseudomonas* species are rarely ingested by *D. discoideum*, we used fluorescence microscopy to visualize interactions between mCherry-labeled QS9 amoebae and GFP-labeled bacteria on agar. After ∼2 d coculture with GFP-labeled *Pseudomonas* and Kp, the amoebae were embedded within a dense lawn of bacteria, but few bacteria were observed within amoebae (Fig. 3C-E). While it is possible to find amoebae that contain individual Pl and Pp cells, intracellular Pv is very uncommon, which is consistent with the results of the gentamicin protection assay. We also imaged mCherry-labeled QS9 amoebae with GFP-labeled *Pseudomonas* and E2crimson-labeled Kp (Fig. 3F-H). Kp is detectable inside of amoebae but, interestingly, large amounts of Kp remain uneaten.

### Some predation-resistant *Pseudomonas* protect edible species

Bacteria may evade predation by secreting proteins or metabolites that suppress predators. If Pv, Pl, or Pp use such a mechanism, we predicted that the presence of predation-resistant bacteria may benefit otherwise edible bacteria in co-culture with *D. discoideum*. We tested this hypothesis with Kp and two edible *Pseudomonas* sp. strains: 14P 8.1_Bac3 (referred to here as Ph) and 7P 10.2_Bac1 (referred to as Pe) (Fig. 4). Pv reduced the effects of predation on all three edible strains, although the effect is only significant for Pe and Ph. An average of 4.12×10^9^ Ph cells were recovered from coculture with Pv, in contrast to 3.76×10^7^ Ph cells recovered in the absence of inedible *Pseudomonas*. Similarly, 6.53×10^9^ Pe cells were recovered from coculture with Pv and 8.33×10 ^7^ Pe cells were recovered from the control. Pp significantly increased the survival of Kp-GFP from 5.78×10^7^ cells without inedible *Pseudomonas* to 2.89×10^9^ cells but reduced the survival of Pe to 2.44×10^3^ cells, suggesting antagonism between *Pseudomonas* strains, though this reduction is not statistically significant. Pl had little effect on the survival of edible bacteria under the conditions tested.

**Figure 4.**
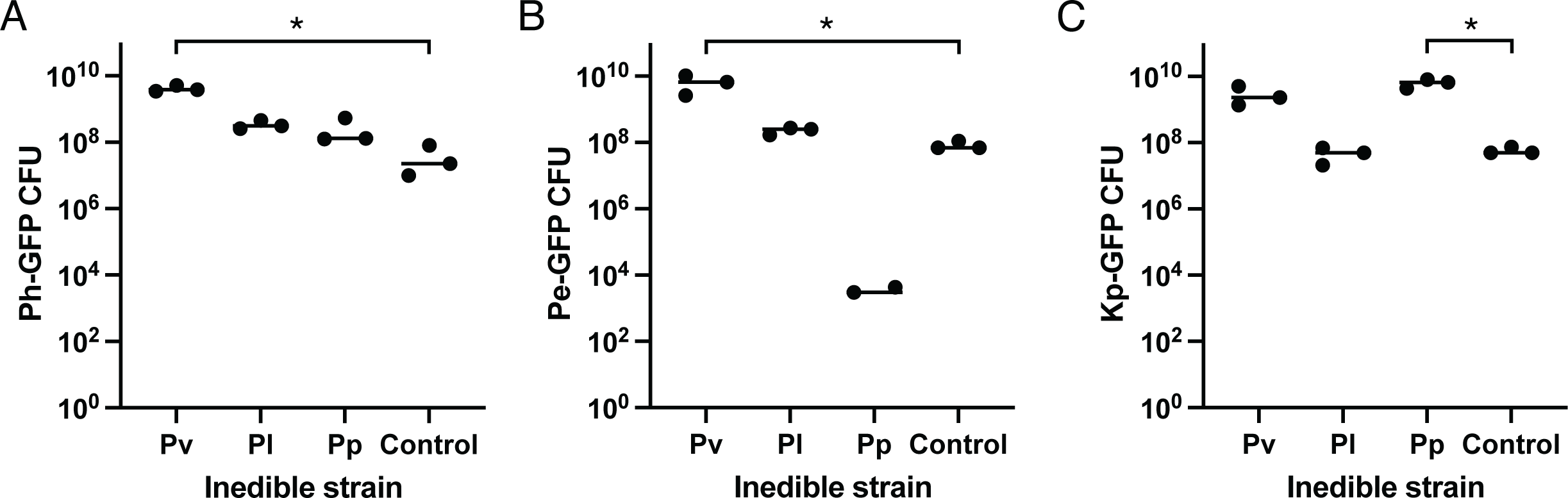
Some inedible *Pseudomonas* strains protect edible bacteria from predation. (A) Ph-GFP, (B) Pe-GFP, and (C) Kp-GFP CFUs recovered after 7 d co-culture with *D. discoideum* AX4. One-way ANOVA with Dunnett’s multiple comparisons test with single pooled variance comparing each sample to the control. *, p ≤ 0.05.

### Diversity of potential predation resistance genes

Several genes and gene clusters are known to help *Pseudomonas* sp. resist predation or survive within the phagosome. These genes include: Type III secretion systems (T3SS); T3SS effectors ExoU and ExoY; Type VI secretion systems (T6SS); ExlA, a pore forming toxin secreted by many *Pseudomonas* species (Basso et al., 2017); and MgtC, which inhibits phagosome acidification and contributes to *Ps. aeruginosa* survival within macrophages (Belon et al., 2015; Garai et al., 2019). To determine whether the distribution of any of these genes could explain why some *Pseudomonas* species are resistant to predation by *D. discoideum*, while others are susceptible, we searched for homologs in the genomes we sequenced as well as related reference genomes (Fig. 5, Table S5). We identified three different T3SS and five different T6SS. A few strains, including Pl, have T3SS and encode homologs to T3SS effectors ExoU and ExoY. While almost all strains encode one or more T6SS, T6SS-3 is present in 5 of 7 predation resistant strains and only 1 of 8 edible strains. Many strains encode homologs of ExlA and MgtC. However, none of these genes or gene clusters are found more frequently in predation resistant strains than edible strains (Fisher’s exact test, p > 0.05, FDR correction for multiple comparisons) (Table S6).

**Figure 5.**
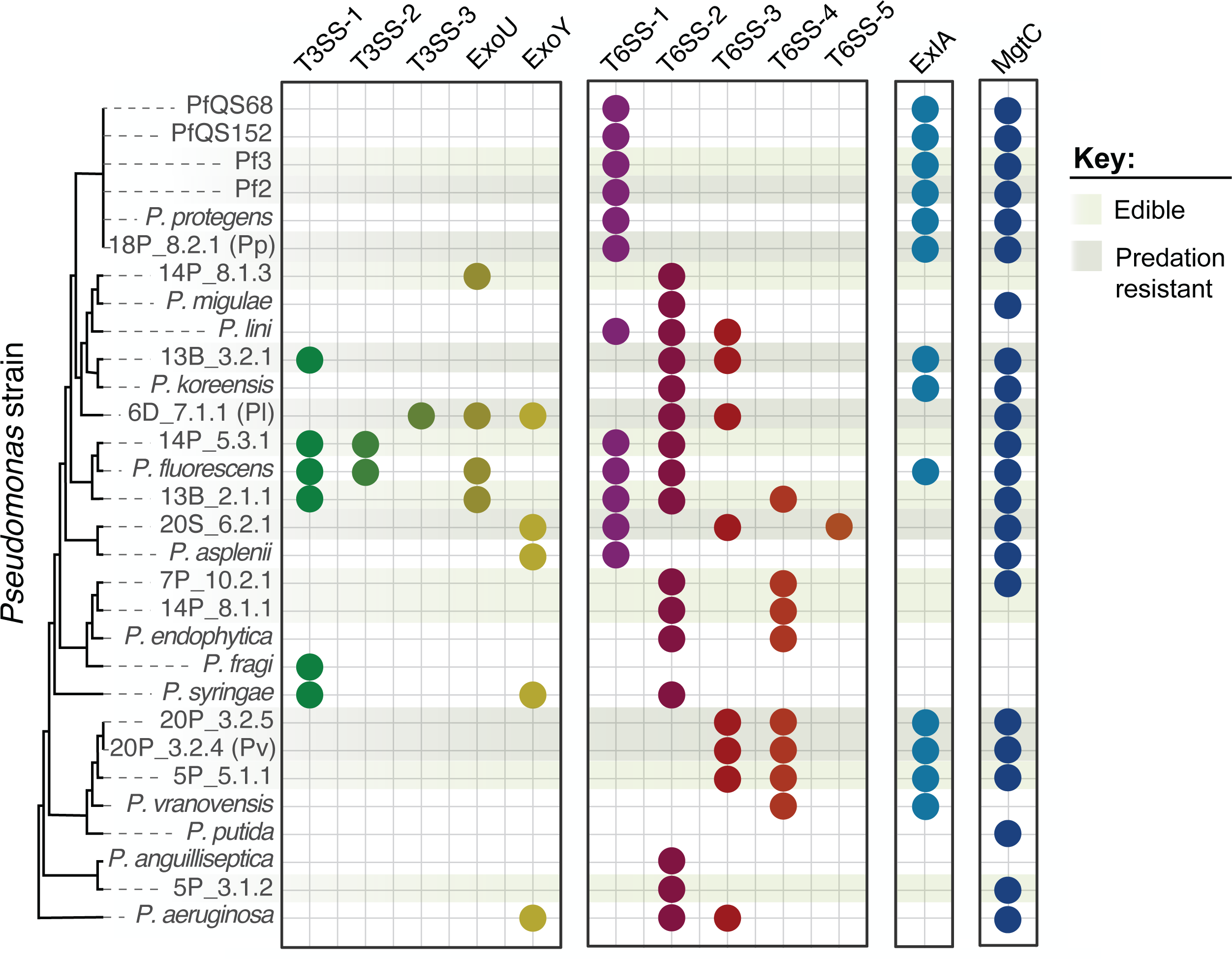
Presence of genes known to contribute to resistance to phagocytosis in other *Pseudomonas* species does not explain predation resistance in soil isolates. Colored circles represent homologs of T3SS structural genes, T3SS effectors ExoU and ExoY, T6SS structural genes, ExlA, and MgtC. Sequenced genomes (identified by isolate name) and reference genomes (identified by species name) are organized based on the whole genome phylogeny from Fig. 1. Light and dark green shading highlights isolates that are edible or resistant to predation.

Since secondary metabolites can also contribute to predation resistance, we used antiSMASH to identify putative secondary metabolite biosynthetic gene clusters, then grouped the gene clusters based on homology (Fig. 6, Fig. S1). Some clusters were found in all or most *Pseudomonas* genomes, while others were limited to certain taxa. None of the secondary metabolite biosynthetic gene clusters are found more frequently in predation resistant strains than in susceptible strains (Fisher’s exact test, p > 0.05, FDR correction for multiple comparisons) (Table S6).

**Figure 6.**
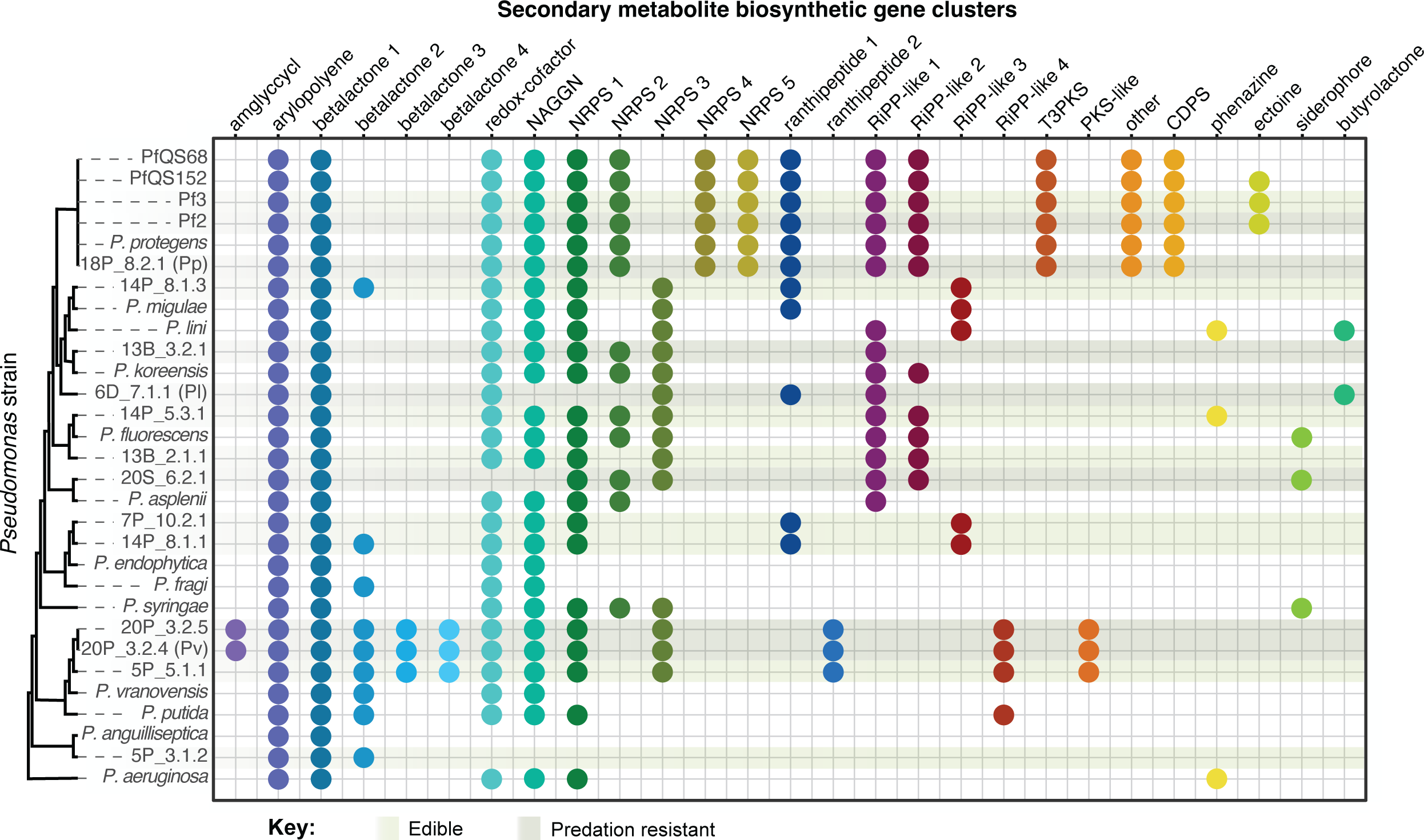
Distribution of secondary metabolite biosynthetic gene clusters in *Pseudomonas* genomes. Colored circles represent the presence of a cluster of biosynthetic genes. Clusters within the same group (x-axis labels) share >70% nucleotide identity over ≥20% of the cluster length. Sequenced genomes (identified by isolate name) and reference genomes (identified by species name) are organized based on the whole genome phylogeny from Fig. 1. Light and dark green shading highlights isolates that are edible or resistant to predation.

## Discussion

Multiple *Pseudomonas* species resist predation by *D. discoideum*, and some infect *D. discoideum* fruiting bodies. However, little is known about most *Pseudomonas* species that have been isolated from *D. discoideum*, many of which appear to belong to novel species based on low ANI with closely related NCBI reference genomes. Pp is a strain of *P. protegens*, a species that has been extensively studied in the context of biocontrol, as *P. protegens* strains produce a variety of secondary metabolites and exoproteins that suppress the growth of fungal and bacterial pathogens of plants (Ramette et al., 2011). Similarly, *P. lini*, a relative of Pl, has been shown to suppress plant pathogens (Gómez-Lama Cabanás et al., 2018). *P. vranovensis*, a relative of Pv, is a pathogen of the nematode *Caenorhabditis elegans* (Burton et al., 2020), but no virulence mechanisms have been described.

Predation resistant strains are not monophyletic, suggesting this trait has been gained or lost multiple times. A few proteins and protein complexes are known to contribute to resistance to phagocytosis or survival within the phagosome in other *Pseudomonas* species. For example, the human pathogen *Ps. aeruginosa* produces the pore-forming toxin Exolysin (ExlA) and secretes exotoxins ExoU and ExoY through a Type III secretion system (T3SS) to kill macrophages (Basso et al., 2017). Among the predation resistant *Pseudomonas* isolates identified in this study, only Pl encoded a T3SS and homologs to ExoU and ExoY. T3SSs were also present in edible strains 14P 5.3_Bac1 and 13B 2.1_Bac1, though these genes were distantly related to the T3SS of Pl and did not share synteny, suggesting the T3SSs are not closely related. *Ps. aeruginosa* also encodes MgtC-like proteins that contribute to virulence and survival within macrophages (Belon et al., 2015) and regulate expression of the T3SS (Garai et al., 2019). In our study, ExlA and MgtC homologs were common among both edible and predation resistant isolates, suggesting that the presence of these genes is not sufficient to confer predation resistance.

Some *P. protegens* strains rely on secondary metabolite production to escape from predation by *D. discoideum* and other protists (Jousset et al., 2006; Stallforth et al., 2013), so we also examined the secondary metabolite biosynthetic gene clusters that are encoded by our *Pseudomonas* isolates. Each predation resistant isolate encoded multiple clusters, with little overlap. The same clusters are found in the predation resistant Pp and Pf2 and edible Pf3 isolates, but Pf3 is known to be edible because of a nonsense mutation in *gacA*, which is part of the two-component system that regulates production of secondary metabolites (Stallforth et al., 2013). Pv shares most of its clusters with the closely related isolate 5P 5.1_Bac1, which is categorized as edible but is not as good a food source as Kp. Pl appears to be missing several clusters that are found in other species, and the clusters it encodes are shared with edible species. Overall, the presence and absence of any one of these genes or gene clusters cannot explain why some *Pseudomonas* strains are edible, while others are predation resistant. The predation resistant strains we identified may use different mechanisms to evade predation. Alternatively, there may be shared predation resistance genes that have not yet been identified or genes that are present in both predation resistant and susceptible strains may be regulated in different ways. Although the mechanism of predation resistance has not yet been identified, the observations that Pv protects edible *Pseudomonas* strains from predation, while Pp protects Kp, are consistent with secretion of anti-predation molecule or protein that can benefit nearby cells. We did not observe a protective effect of Pl, which may mean that it does not secrete predation resistance molecules or that the production of such molecules is temperature dependent, like its ability to infect the sorus. Convergence on a predation-resistant phenotype, even if it is through different mechanisms, emphasizes the influence of predation on the evolution of bacteria in soil communities.

Based on our microscopy and gentamicin protection assays, Pv, Pl, and Pp appear to be ingested less frequently than edible Kp and symbiotic *Pa. bonniea*. We recovered substantial amounts of Kp from amoebae lysed a few hours after antibiotic treatment, which is consistent with results of studies that use similar assays (Pukatzki et al., 2002). In our experiments, *D. discoideum* was able to digest the small amounts of Pv and Pl tha t it consumed, suggesting that the mechanism of predation resistance in these two species likely depends on not being ingested. Though Pp was able to persist after phagocytosis, the number of intracellular bacteria is small and decreases over time, suggesting Pp does not replicate intracellularly.

Interestingly, we found that at least one *Pseudomonas* species (Pv) can induce secondary infections of *D. discoideum* fruiting bodies by otherwise edible bacteria. This trait has previously been associated with three *Paraburkholderia* species that are intracellular symbionts of *D. discoideum* (DiSalvo et al., 2015; Haselkorn et al., 2019; Khojandi et al., 2019). By inducing *D. discoideum* to carry or “farm” edible bacteria, symbiotic *Paraburkholderia* are beneficial to the host when spores disperse to areas where prey bacteria are scarce, even though they reduce spore production (Scott et al., 2022). These *Paraburkholderia* species are intracellular and efficiently infect spores, allowing them to disperse with spores and remain associated with *D. discoideum* over many generations. As a result, they likely experience selective pressure to minimize negative effects on the host, as vertically transmitted symbionts may be more likely to become mutualists rather than parasites (Sachs et al., 2011). *Pa. hayleyella* and *Pa. bonniea* demonstrate genome reduction (Brock et al., 2020), increased proportions of infected spores, and reduced numbers of bacteria per spore (Miller et al., 2020) when compared to *Pa. agricolaris*, which is consistent with adaptation to a symbiotic lifestyle. In contrast, *Pseudomonas* infections of fruiting bodies appear to be strictly extracellular and only some fruiting bodies become infected, which would naturally lead to a much less stable association between the bacteria and the host. However, bacteria that infect the sorus could still co-disperse with spores, potentially leading to selection for mutualistic or parasitic traits. Purely opportunistic interactions between bacteria and amoebae may be the evolutionary origin of more complex symbioses.

In conclusion, multiple environmental *Pseudomonas* species evade predation by *D. discoideum* and exhibit some traits characteristic of the *D. discoideum-Paraburkholderia* symbiosis. These symbiont-like behaviors, including infecting the sorus, persisting inside of amoebae, and inducing secondary infections, are likely byproducts of the mechanisms these bacteria use to promote their own survival when interacting with *D. discoideum* and other predators, rather than symbiotic adaptations. However, the presence of these traits in environmental bacteria suggests that the threshold for establishing symbiosis with *D. discoideum* may be low. The genes responsible for predation resistance in Pv, Pl, and Pp have not yet been identified, but it seems likely that predation resistance has evolved multiple times within *Pseudomonas* and may be achieved by multiple means.

## Data availability

16S rRNA gene sequences and genomes are available through Genbank (ON954494-ON954502 and JANLNW000000000-JANLOH000000000), while raw Illumina reads are available through the NCBI SRA (PRJNA857029). Code is available at https://github.com/misteele/Dicty-Pseudomonas-genomes.

## Acknowledgements

We thank the Dictybase Stock Center for providing *D. discoideum* AX4 and Kp, Julie Perreau, Nancy Moran, Jeffrey Barrick, and Susanne DiSalvo for providing plasmids, Debbie Brock for protocols and support, Shreenidhi PM for providing the *D. discoideum* QS9-mCherry clone, and members of the Queller/Strassmann lab for useful discussions.

This work is based upon work supported by the National Science Foundation (DEB 1753743 and DEB2237266 to JES/DCQ and NSF Postdoctoral Research Fellowships in Biology Program Grant No. 2109487 to MIS).

