## Supplementary material for "Predation-resistant *Pseudomonas* bacteria engage in symbiont-like behavior with the social amoeba *Dictyostelium discoideum*"

**Table S1. Strains and plasmids used in this study.**

| Species | Strain | Abbreviation | Description | Source |
|---|---|---|---|---|
| *Escherichia coli* | MFD*pir* | | Donor strain for conjugations | [1] |
| | WM3064 | | Donor strain for conjugations | [2] |
| *Klebsiella pneumoniae* | Kp | | | (Dictybase stock center) |
| | Kp-Gm-GFP | | | (this study) |
| | Kp-Gm-E2crimson | | | (this study) |
| *Paraburkholderia bonniea* | Bb859 | Bb | | |
| | Bb859-Gm-GFP | Bb-GFP | | (this study) |
| *Pseudomonas aeruginosa* | PAO1 | Pa | | |
| | PAO1-GFP | Pa-GFP | | [3] |
| *Pseudomonas* sp. | 5P_3.1_Bac2 | | Isolate | [4] |
| | 5P_5.1_Bac1 | | Isolate | [4] |
| | 5P_5.1_Bac1-Km-GFP | | Chromosomal insertion of KmR and GFP at attTn7 site | (this study) |
| | 20P_3.2_Bac4 | Pv | Isolate | [4] |
| | 20P_3.2_Bac4-Km-GFP | Pv-GFP | Chromosomal insertion of KmR and GFP at attTn7 site | (this study) |
| | 20P_3.2_Bac5 | | Isolate | [4] |
| | 20P_3.2_Bac4-Gm-GFP | | Chromosomal insertion of GmR and GFP at attTn7 site | (this study) |
| | 14P_8.1_Bac1 | | Isolate | [4] |
| | 7P_10.2_Bac1 | Pe | Isolate | [4] |
| | 7P_10.2_Bac1-Km-GFP | Pe-GFP | Chromosomal insertion of KmR and GFP at attTn7 site | (this study) |
| | 20S_6.2_Bac1 | | Isolate | [4] |
| | 20S_6.2_Bac1-Gm-GFP | | Chromosomal insertion of GmR and GFP at attTn7 site | (this study) |
| | 13B_2.1_Bac1 | | Isolate | [4] |
| | 14P_5.3_Bac1 | | Isolate | [4] |

| | | | | |
|---|---|---|---|---|
| | 14P_5.3_Bac1-Gm-GFP | | Chromosomal insertion of GmR and GFP at attTn7 site | (this study) |
| | 6D_7.1_Bac1 | Pl | Isolate | [4] |
| | 6D_7.1_Bac1-Gm-GFP | Pl-GFP | Chromosomal insertion of GmR and GFP at attTn7 site | (this study) |
| | 13B_3.2_Bac1 | | Isolate | [4] |
| | 13B_3.2_Bac1-Gm-GFP | | Chromosomal insertion of GmR and GFP at attTn7 site | (this study) |
| | 14P_8.1_Bac3 | Ph | Isolate | [4] |
| | 14P_8.1_Bac3-Gm-GFP | Ph-GFP | Chromosomal insertion of GmR and GFP at attTn7 site | (this study) |
| *Pseudomonas protegens* | 18P_8.2_Bac1 | Pp | Isolate | [4] |
| | 18P_8.2_Bac1-Gm-GFP | Pp-GFP | Chromosomal insertion of GmR and GFP at attTn7 site | (this study) |
| | Pf2 | | Isolate | [5] |
| | Pf3 | | Isolate | [5] |
| *Dictyostelium discoideum* | AX4 | | Axenic clone | (Dictybase stock center) |
| | QS157 | | Wild clone | |
| | QS9-mCherry | | Wild clone with chromosomal mCherry | |

| | Plasmid | Description | Source |
|---|---|---|---|
| | pTNS2 | Tn7 transposase | [6] |
| | pUC18R6KT-miniTn7T-Gm-sfGFP | mini-Tn7 transposon carrying GmR and GFP | [7] |
| | pUC16R6KT-miniTn7-Gm-PA3-E2crimson | mini-Tn7 transposon carrying GmR and E2crimson | Julie Perreau |
| | pURR25 | mini-Tn7 transposon carrying KmR and GFP | [2] |
| | pUX-BF13 | Tn7 transposase | [2] |

**Table S2.** The number of infected and uninfected sori collected from *D. discoideum* fruiting bodies grown on mixtures of *Pseudomonas* sp. and Kp. Fisher's exact test was used to compare the number of infected and uninfected sori for each strain to the total number of infected and uninfected sori. False discovery rate (FDR) was used to correct p values for multiple comparisons.

| Strain | Strain edibility | Infected sori | Uninfected sori | FDR adjusted pvalue |
|---|---|---|---|---|
| **Pf3** | Edible | 0 | 60 | 2.14E-12 |
| **14P_8.1_Bac3 (Ph)** | Edible | 1 | 49 | 6.42E-09 |
| **13B_2.1_Bac1** | Edible | 0 | 50 | 1.89E-10 |
| **7P_10.2_Bac1 (Pe)** | Edible | 5 | 45 | 2.47E-05 |
| **14P_8.1_Bac1** | Edible | 10 | 30 | 1.11E-01 |
| **5P_5.1_Bac1** | Edible | 36 | 34 | 7.31E-02 |
| **5P_3.1_Bac2** | Edible | 7 | 53 | 1.27E-05 |
| **14P_5.3_Bac1** | Edible | 5 | 5 | 5.28E-01 |
| **Pf2** | Inedible | 13 | 7 | 5.35E-02 |
| **18P_8.2_Bac1 (Pp)** | Inedible | 18 | 2 | 1.27E-05 |
| **6D_7.1_Bac1 (Pl)** | Inedible | 34 | 31 | 6.78E-02 |
| **20S_6.2_Bac1** | Inedible | 1 | 9 | 1.11E-01 |
| **20P_3.2_Bac4 (Pv)** | Inedible | 75 | 5 | 1.66E-21 |
| **20P_3.2_Bac5** | Inedible | 68 | 9 | 1.74E-16 |
| **13B_3.2_Bac1** | Inedible | 2 | 8 | 3.51E-01 |
| **PAO1** | Inedible | 4 | 32 | 5.91E-04 |

**Table S3.** Temperature affects the fraction of *D. discoideum* sori that become infected with Pl.

No fruiting bodies developed on 90% Pl plates incubated at 18ºC.

| | | 18ºC incubation | | 25ºC incubation | |
|---|---|---|---|---|---|
| Treatment | Replicate | Spots with bacteria | Total spots | Spots with bacteria | Total spots |
| 10% Pl | 1 | 5 | 10 | 0 | 10 |
| | 2 | 5 | 10 | 0 | 10 |
| 50% Pl | 1 | 2 | 5 | 0 | 3 |
| | 2 | 5 | 8 | 0 | 10 |
| 90% Pl | 1 | | 0 | 0 | 1 |
| | 2 | | 0 | 0 | 3 |

**Table S4.** Average Nucleotide Identity between isolate genomes and the NCBI reference genome with the most similar 16S rRNA sequence. Pp is closely related to *Pseudomonas protegens* CHA0 (98.8% ANI). Pv and 20P 3.2_Bac5, which are nearly identical (99.99% ANI), are most closely related to *Pseudomonas alkylphenolica* (84.5 and 84.6% ANI) but were identified as *Pseudomonas vranovensis* (84.3 and 84.4% ANI) when they were first isolated. Pl was originally identified as *Pseudomonas lini* (85.7% ANI) but is more closely related to *Pseudomonas frederiksbergensis* (85.9% ANI).

| Isolate | Abb. | Reference genome | Reference accession | ANI |
|---|---|---|---|---|
| 13B 2.1_Bac1 | | *Pseudomonas lurida* | GCF_001708485.1 | 89.08 |
| 14P 5.3_Bac1 | | *Pseudomonas fluorescens* | GCF_900215245.1 | 89.51 |
| 14P 8.1_Bac1 | | *Pseudomonas helleri* | GCF_001043025.1 | 96.48 |
| 14P 8.1_Bac3 | Ph | *Pseudomonas migulae* | GCF_900106025.1 | 89.37 |
| 20P 3.2_Bac4 | Pv | *Pseudomonas alkylphenolica* | GCF_009755645.1 | 84.52 |
| 20P 3.2_Bac5 | | *Pseudomonas alkylphenolica* | GCF_009755645.1 | 84.61 |
| 20S 6.2_Bac1 | | *Pseudomonas asplenii* | GCF_900105475.1 | 85.18 |
| 5P 3.1_Bac2 | | *Pseudomonas anguilliseptica* | GCF_900105355.1 | 81.48 |
| 6D 7.1_Bac1 | Pl | *Pseudomonas frederiksbergensis* | GCF_002967995.1 | 85.88 |
| 7P 10.2_Bac1 | Pe | *Pseudomonas helleri* | GCF_001043025.1 | 89.03 |
| 18P 8.2_Bac1 | Pp | *Pseudomonas protegens* | GCF_900560965.1 | 98.77 |
| Pf2 | | *Pseudomonas protegens* | GCF_900560965.1 | 98.77 |
| Pf3 | | *Pseudomonas protegens* | GCF_900560965.1 | 98.73 |
| PfQS152 | | *Pseudomonas protegens* | GCF_900560965.1 | 98.77 |
| PfQS68 | | *Pseudomonas protegens* | GCF_900560965.1 | 98.82 |

**Table S5.** Amino acid percent identity shared between ExlA, ExoU, ExoY, and MgtC homologs identified in *Pseudomonas* isolate genomes and *Ps. aeruginosa* reference sequences.

| Strain | ExlA | ExoU | ExoY | MgtC |
|---|---|---|---|---|
| *Pseudomonas aeruginosa* | | | 99.74 | 100 |
| *Pseudomonas* sp. 5P_3_1_Bac2 | | | | 46.52 |
| *Pseudomonas anguilliseptica* | | | | |
| *Pseudomonas putida* | | | | 86.92 |
| *Pseudomonas vranovensis* | 39.16 | | | |
| *Pseudomonas* sp. 5P_5_1_Bac1 | 39.73 | | | 87.61 |
| *Pseudomonas* sp. 20P_3_2_Bac4 (Pv) | 39.51 | | | 88.03 |
| *Pseudomonas* sp. 20P_3_2_Bac5 | 39.51 | | | 88.03 |
| *Pseudomonas syringae* | | | 24.61 | |
| *Pseudomonas fragi* | 51.42 | | | |
| *Pseudomonas endophytica* | | | | |
| *Pseudomonas* sp. 14P_8_1_Bac1 | | | | |
| *Pseudomonas* sp. 7P_10_2_Bac1 (Pe) | | | | 44.2 |
| *Pseudomonas asplenii* | | | 27.93 | 45.25 |
| *Pseudomonas* sp. 20S_6_2_Bac1 | | | 30.58 | 44.55 |
| *Pseudomonas* sp. 13B_2_1_Bac1 | 56.65 | 43.36 | | 43.81 |
| *Pseudomonas fluorescens* | 59.4 | 45.46 | | 44.25 |
| *Pseudomonas* sp. 14P_5_3_Bac1 | | | | 43.81 |
| *Pseudomonas* sp. 6D_7_1_Bac1 (Pl) | | 62.94 | 33.69 | 43.11 |
| *Pseudomonas koreensis* | 56.5 | | | 44 |
| *Pseudomonas* sp. 13B_3_2_Bac1 | | | | 44 |
| *Pseudomonas lini* | | | | |
| *Pseudomonas migulae* | | | | 44.89 |
| *Pseudomonas* sp. 14P_8_1_Bac3 (Ph) | | 45.19 | | |
| *Pseudomonas protegens* 18P_8_2_Bac1 (Pp) | 59.35 | | | 44.25 |
| *Pseudomonas protegens* | 59.41 | | | 44.25 |
| *Pseusomonas fluorescens* Pf2 | 59.35 | | | 44.25 |
| *Pseusomonas fluorescens* Pf3 | 59.35 | | | 44.25 |
| *Pseusomonas fluorescens* PfQS152 | 59.35 | | | 44.25 |
| *Pseusomonas fluorescens* PfQS68 | 59.27 | | | 44.69 |

**Table S6.** For each putative predation resistance gene, Fisher's exact test was used to compare the number of edible strains with and without the gene to the number of inedible strains with and without the gene. FDR corrected p values are shown.

| Gene(s) | Edible strains with gene | Edible strains without gene | Inedible strains with gene | Inedible strains without gene | p value | FDR corrected p value |
|---|---|---|---|---|---|---|
| T3SS-1 | 2 | 6 | 1 | 6 | 1.00 | 1.00 |
| T3SS-2 | 1 | 7 | 0 | 7 | 1.00 | 1.00 |
| T3SS-3 | 0 | 8 | 1 | 6 | 0.47 | 0.83 |
| ExoU | 2 | 6 | 1 | 6 | 1.00 | 1.00 |
| ExoY | 0 | 8 | 2 | 5 | 0.20 | 0.83 |
| T6SS-1 | 3 | 5 | 3 | 4 | 1.00 | 1.00 |
| T6SS-2 | 6 | 2 | 2 | 5 | 0.13 | 0.83 |
| T6SS-3 | 1 | 7 | 5 | 2 | 0.04 | 0.83 |
| T6SS-4 | 4 | 4 | 2 | 5 | 0.61 | 0.83 |
| T6SS-5 | 0 | 8 | 1 | 6 | 0.47 | 0.83 |
| ExlA | 2 | 6 | 5 | 2 | 0.13 | 0.83 |
| MgtC | 6 | 2 | 7 | 0 | 0.47 | 0.83 |
| Amglyccycl | 0 | 8 | 2 | 5 | 0.20 | 0.83 |
| Betalactone 1 | 4 | 4 | 2 | 5 | 0.61 | 0.83 |
| Betalactone 3 | 1 | 7 | 2 | 5 | 0.57 | 0.83 |
| Betalactone 4 | 1 | 7 | 2 | 5 | 0.57 | 0.83 |
| Redox-cofactor | 7 | 1 | 6 | 1 | 1.00 | 1.00 |
| NAGGN | 7 | 1 | 5 | 2 | 0.57 | 0.83 |
| NRPS 1 | 7 | 1 | 6 | 1 | 1.00 | 1.00 |
| NRPS 2 | 2 | 6 | 4 | 3 | 0.31 | 0.83 |
| NRPS-like 3 | 4 | 4 | 5 | 2 | 0.61 | 0.83 |
| NRPS 4 | 1 | 7 | 2 | 5 | 0.57 | 0.83 |
| NRPS 5 | 1 | 7 | 2 | 5 | 0.57 | 0.83 |
| Ranthipeptide 1 | 4 | 4 | 3 | 4 | 1.00 | 1.00 |
| Ranthipeptide 2 | 1 | 7 | 2 | 5 | 0.57 | 0.83 |
| RiPP-like 1 | 3 | 5 | 5 | 2 | 0.31 | 0.83 |
| RiPP-like 2 | 3 | 5 | 3 | 4 | 1.00 | 1.00 |
| RiPP-like 3 | 3 | 5 | 0 | 7 | 0.20 | 0.83 |
| RiPP-like 4 | 1 | 7 | 2 | 5 | 0.57 | 0.83 |
| T3PKS | 1 | 7 | 2 | 5 | 0.57 | 0.83 |
| PKS-like | 1 | 7 | 2 | 5 | 0.57 | 0.83 |
| Other | 1 | 7 | 2 | 5 | 0.57 | 0.83 |

| | | | | | | |
|---|---|---|---|---|---|---|
| CDPS | 1 | 7 | 2 | 5 | 0.57 | 0.83 |
| Phenazine | 1 | 7 | 0 | 7 | 1.00 | 1.00 |
| Ectoine | 1 | 7 | 1 | 6 | 1.00 | 1.00 |
| Siderophore | 0 | 8 | 1 | 6 | 0.47 | 0.83 |
| Butyrolactone | 0 | 8 | 1 | 6 | 0.47 | 0.83 |

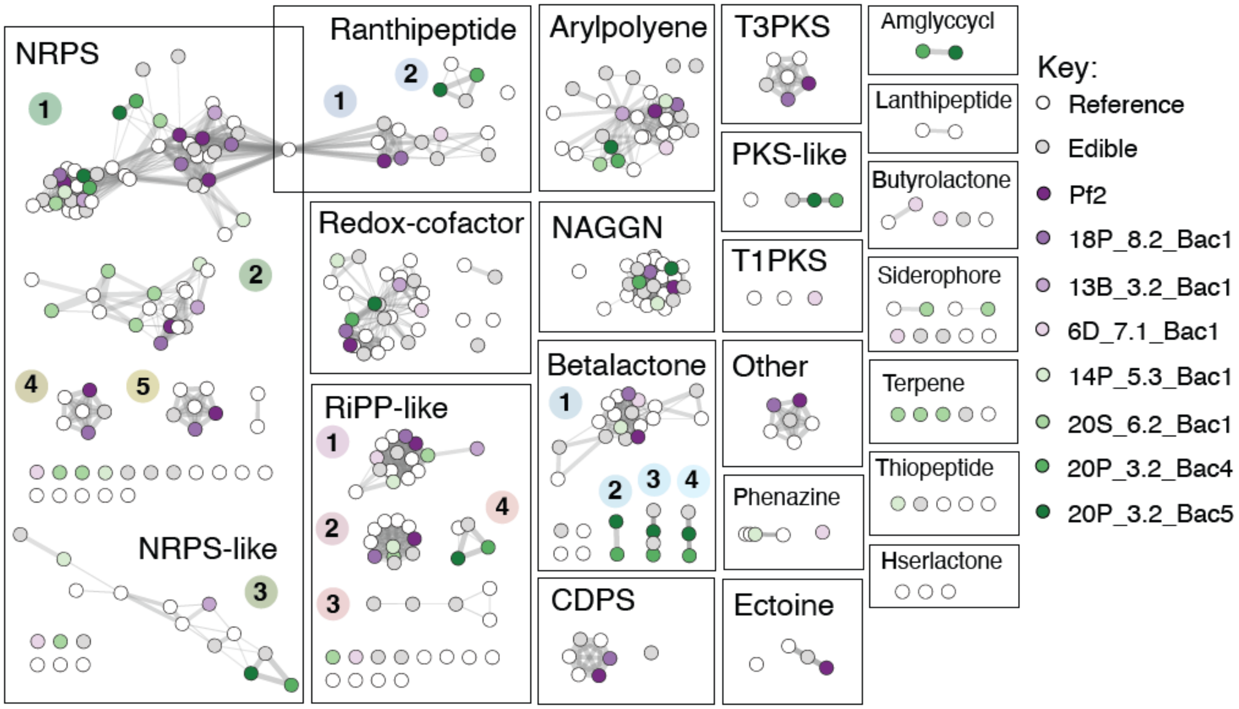


**Figure S1. No secondary metabolite biosynthetic gene clusters are unique to predation resistant Pseudomonas genomes.** Homology network diagram showing relationships between secondary metabolite gene clusters in *Pseudomonas* genomes. Nodes represent nucleotide sequences of gene clusters. Nodes representing sequences from predation resistant isolates are colored according to species. Grey nodes represent sequences from edible isolates, while white nodes are sequences from reference genomes. Lines connect nodes that share >70% nucleotide identity over more than 20% of the total length of the cluster. Line color represents percent identity, while width is proportional to alignment length relative to query length. Clusters are numbered when more than one cluster of a kind was identified.
